# PLK4 inhibition as a strategy to enhance non-small cell lung cancer radiosensitivity

**DOI:** 10.1101/2025.02.19.638860

**Authors:** Irma G. Dominguez-Vigil, Kishore Banik, Marta Baro, Joseph N. Contessa, Thomas J. Hayman

**Author notes:** **Corresponding author:** Thomas J. Hayman; 15 York Street, New Haven, CT 06510. These authors contributed equally (Irma G. Dominguez-Vigil, Kishore Banik).

## Abstract

Lung cancer is the leading cause of cancer-related mortality worldwide. Non-small cell lung cancer (NSCLC) is the most common subtype of lung cancer and comprises 85% of cases. Despite treatment advances, local control after curative-intent chemoradiation for NSCLC remains suboptimal. Polo-like kinase 4 (PLK4) is a serine-threonine kinase that plays a critical role in the regulation of centrosome duplication and cell cycle progression and is overexpressed in NSCLC, thus, making it a potential therapeutic target. CFI-400945 is an orally available PLK4 inhibitor currently undergoing clinical trial evaluation. As radiation causes cell death primarily by mitotic catastrophe, a process enhanced by alterations in centrosome amplification, we hypothesized that disruption of the mitotic machinery by inhibition of PLK4 would enhance the effects of radiation in NSCLC. PLK4 inhibition by CFI-400945 resulted in radiosensitization of NSCLC cell lines. In contrast, CFI-400945 had no effect on the radiosensitivity of normal lung fibroblasts. PLK4 inhibition did not affect cell-cycle phase distribution prior to radiation, but rather the combination of CFI-400945 and radiation resulted in increased G2/M cell cycle arrest, increased centrosome amplification, and a concomitant increase in cell death through mitotic catastrophe. Lastly, CFI-400945 treatment enhanced the radiation-induced tumor growth delay of NSCLC tumor xenografts. These data indicate that targeting PLK4 is a novel approach to enhance the radiation sensitivity of NSCLC *in vitro* and *in vivo* through potentiation of centrosome amplification and cell death through mitotic catastrophe.

## Introduction

Lung cancer affects greater than 2 million patients annually worldwide and remains the leading cause of cancer-related deaths (1). Non-small cell lung cancer (NSCLC), accounts for approximately 85% of lung cancer diagnoses (2). Radiation therapy is a primary curative treatment modality for patients with both early stage and locally advanced NSCLC (3,4). However, despite treatment intensification, outcomes remain poor in the locally advanced setting with up to 40% of patients with locally advanced NSCLC experiencing locoregional recurrence (5). As such previous efforts have been focused on improving standard chemoradiation treatment as a means to increase disease control and ultimately patient survival. Towards this end, the PACIFIC trial demonstrated improvement in 4-year overall survival (OS) and progression-free survival (PFS) to 50% and 35% respectively with the addition of adjuvant durvalumab to chemoradiation (6,7). Among these failures, local-only failure occurred in ∼36% of patients in the PACIFIC trial (8). While these findings represent a significant improvement in NSCLC treatment, they still indicate that ∼65% of patients ultimately develop progressive disease, with a substantial portion occurring as local only failures, indicating the need for novel therapies to improve curative intent chemoradiation.

Maintenance of genomic stability after genotoxic stress such as radiation is critical to avoid cell death (9,10). Centrosomes, which are small membrane-free organelles composed of two centrioles surrounded by a pericentriolar matrix, are a critical determinant of genomic stability (11). Centrosome duplication occurs once per cell cycle in a tightly regulated manner to ensure proper chromosome segregation (11). Processes leading to dysregulation of centriole duplication will cause centrosome amplification, defined as the presence of >2 centrosomes (12–15). In the context of DNA damage, centrosome amplification has been shown to result in increased cell death through mitotic catastrophe (16–19). Thus, strategies to enhance tumor centrosome amplification may potentially be exploited for therapeutic gain in combination with DNA damaging therapies such as radiation.

The Polo-like kinases (PLKs) are a family of serine-threonine kinases compromised of five family members, PLK1-5 (20,21). The PLKs regulate a diverse array of cellular processes including cell cycle progression, mitosis, cytokinesis, and centrosome function (20,21). PLK4 is a structurally unique member of this family that has a central role in the regulation of centriole and centrosome duplication through its interaction with CEP (centriolar protein) family members and other proteins (22–24). Overexpression of PLK4 has been demonstrated in NSCLC and other malignancies and this overexpression is associated with worse OS, PFS, and more advanced disease (25–27). Furthermore, in preclinical models, aberrant expression of PLK4 leads to centrosome amplification and chromosomal instability culminating in tumorigenesis (28–31).

These findings have led to significant interest in pharmacologic targeting of PLK4 as a therapeutic strategy to improve outcomes in multiple malignancies. CFI-400945 is a first-in-class, orally available PLK4 inhibitor that is currently undergoing clinical trial evaluation in multiple tumor types (24–26,29,32,33). It is an ATP-competitive kinase inhibitor that how been shown to selectively inhibit PLK4 at low nanomolar concentrations (IC50 2.8nM for PLK4 vs 98nM for Aurora Kinase B) (33). At low nanomolar concentrations, CFI-400945 has been shown to partially inhibit PLK4 activity resulting in centrosome amplification (similar to other preclinical PLK4 inhibitors) (26,33–35). CFI-400945 treatment impairs proliferation, induces polyploidy, and increases cell death in several preclinical tumor models (25,26,33,35). Clinically the results of a Phase 1 dose-escalation trial in patients with advanced solid tumors have shown that CFI-400945 is well-tolerated and have established a Phase 2 dose (36,37). Given the critical role of PLK4 in controlling centrosome amplification, coupled with the importance of centrosome amplification in the response to DNA damaging therapies we sought to investigate the effects of PLK4 inhibition using CFI-400945 on the radiation response of NSCLC. Our results indicate that targeting PLK4 is a novel approach to enhance the radiation sensitivity of NSCLC through potentiation of centrosome amplification and cell death through mitotic catastrophe.

## Methods

### Cell lines and treatment

The human NSCLC cell lines H460 and A549, and human normal lung fibroblast cell line MRC9 were purchased from American Type Culture Collection (ATCC). H460 and A549 were cultured in RPMI 1640 (Gibco, Life Technologies), supplemented with 10% Fetal Bovine Serum (Gibco, Life Technologies). MRC9 cells were cultured in Advanced MEM media (Gibco) supplemented with 10% Fetal Bovine Serum (Gibco). Cells were cultured in a humidified incubator with 5% CO_2_ and kept in culture no more than two months after resuscitation from the original stocks. Mycoplasma cell culture contamination was ruled out using MycoAlert Mycoplasma Detection Kit (Lonza) or Universal Mycoplasma Detection Kit (ATCC).

Cell cultures were irradiated using a Precision X-Ray 320 kV voltage unit at a dose rate of 2.3 Gy/min or 2.6Gy/min with a 2-mm aluminum filter (Precision X-Ray Inc, USA) with the dose indicated in the respective figure legend. Quality assurance was performed monthly using a P.T.W. 0.3 cm^3^ ionization chamber calibrated to NIST standards and quarterly dosimetry using thermoluminescent dosimeter-based or ferrous sulfate-based dosimeters. Dose-mapping is performed by Precision X-Ray annually. Cells were treated with CFI-400945 (kindly provided by Dr. Tak Mak or purchased through Selleckchem) or Centrinone (MedChemExpress) dissolved in dimethyl sulfoxide (DMSO). DMSO was used as a vehicle control.

### Cell proliferation assay

Cell proliferation was determined using the MTT (3-[4,5-dimethylthiazol-2-yl]-2,5 diphenyl tetrazolium bromide) assay. Specifically, two thousand A549 or H460 cells were seeded in triplicates in 96-well plates and incubated for 24 hours. Subsequently, the cells were treated with varying concentrations of CFI-400945 and Centrinone as indicated. Cell numbers were estimated by adding MTT reagent to the wells at the indicated times and spectrophotometric readings (absorbance at 570nM) were obtained after 3h. Control cells were considered to have 100% proliferation, and the percentage proliferation was calculated using the following formula: (Absorption of treated cells × 100) / (Absorption of control cells (vehicle)).

### Clonogenic survival assay

Cells were plated, allowed to attach overnight, and then treated with the indicated dose of CFI-400945 or Centrinone for 24h. Cells were then irradiated as indicated and 24h after RT cells were washed, trypsinized, and plated at clonal density in six-well plates to determine clonogenic survival. Subsequently, 10-14 days after seeding, the plates were stained with 0.25% crystal violet in 80% methanol. Colonies containing more than 50 cells (H460 and A549), or 25 cells (MRC9) were then counted. The surviving fraction of each sample was determined by calculating the ratio of the number of colonies counted to the number of cells seeded, normalized for plating efficiency differences due to treatment. Differences in clonogenic survival for each treatment were assessed by comparing survival curves generated from the linear quadratic equation as previously described (38).

### Immunoblotting assay

Immunoblotting was performed as previously described (39). Nitrocellulose-bound primary antibodies were detected with anti-rabbit IgG horseradish peroxidase-linked antibody (Millipore-Sigma) and bands were visualized with Amersham ECL detection reagent (Cytiva). Anti-PLK4 antibody was obtained from Thermo Scientific (# PA529373, 1:1,000) and anti-β-tubulin was obtained from Cell Signaling Technologies, (#2128, 1:5,000).

### Cell cycle and DNA content analysis

Cell-cycle distribution was determined by flow-cytometry based analysis of DNA content. Briefly cells were treated as indicated and fixed in 70% ethanol, stained with FxCycle Propidium Iodide (PI)/RNase Staining Solution (Invitrogen) analyzed using a LSR II flow cytometer (BD Biosciences). At least 10,000 cells per condition were analyzed with data shown representing the mean and SEM of three independent experiments. Double thymidine block was used for synchronization as indicated. Specifically, 2 mM thymidine (Sigma-Aldrich) was first added to cells for 18 h, withdrawn for 9 h, and blocked again with thymidine 2 mM for another 16 h. Cells were treated as indicated and collected for cell cycle analysis as described above.

### Centrosome immunofluorescence analysis

Cells were grown on coverslips and after the indicated treatment, cells were fixed in cold Methanol and permeabilized with 0.2% Triton X-100 in PBS and blocked in 10% goat serum (Sigma) overnight, followed by an overnight incubation with Anti-y-Tubulin (Millipore, #T6557) primary antibody at 1:500 dilution at 4°C. Fixed cells were then incubated with Alexa Fluor 488 goat anti-mouse (Invitrogen) secondary antibody at 1:500 dilution for 1.5 hours at room temperature. Nuclei were stained with DAPI-Hoechst 33342. Coverslips were mounted in Dako Fluorescence Mounting Medium (Agilent) and imaged on a Stellaris 8 Falcon Confocal Microscope (Leica) at 63x for analysis or 100x for representative images. Data presented are mean ± SEM 3 biological replicates with ≥50 cells per replicate counted for each experimental condition.

### Markers of mitotic catastrophe

Cells were grown in chamber slides (Thermo Scientific). After the indicated treatment, cells were fixed in 4% neutral buffered formaldehyde, permeabilized with 0.1% Triton X-100, and blocked with 1% bovine serum albumin in PBS-tween containing 5% goat serum. Slides were then incubated overnight with anti-α-tubulin (Sigma-Aldrich, # T5618) followed by incubation with goat anti-mouse with Alexa-488 secondary antibody (Invitrogen, # A11029) both at 1:1000 dilution. Slides were mounted with Prolong gold anti-fade reagent (Invitrogen) containing 4’, 6-diamidino-2-phenylindole (DAPI) and imaged on a Leica SP5 confocal microscope or EVOS M5000 Fluorescent Microscope for analysis. At least 75 cells per replicate were counted for each experimental condition. Cells with nuclear fragmentation, defined as the presence of two or more distinct nuclear lobes within a single cell were classified as being in mitotic catastrophe as previously reported (17,40–43).

### *In vivo* tumor growth delay

H460 and A549 tumor xenografts were established by injection of 5 × 10^6^ cells subcutaneously into the right hind leg of 4-6-week-old female athymic nude mice (*Foxn1^nu^*, Envigo). When tumor xenografts reached ∼200 mm^3^ volume, animals were randomized into four groups: control, CFI-400945, RT, and the combination of CFI-400945 plus RT. For the control group, mice were treated with vehicle (sterile water, Invitrogen) daily by oral gavage for a total of 6 days. For the CFI-400945 group, mice were treated with 7.5 mg/Kg of CFI400945 daily dose by oral gavage for a total of 6 days. For the RT group, fractionated radiation was delivered locally using XRAD-320 at a dose rate of 3.3 Gy/min with a 2-mm aluminum filter (Precision X-Ray Inc) at 2 Gy daily for 5 days. For the combination treatment group, mice received CFI-400945 (7.5 mg/kg daily) for a total of 6 days, starting the first day of radiation. Tumor size was measured three times a week and calculated according to the formula (L × W^2^)/2. Tumor growth delay was calculated as the time to reach 400 mm^3^, representing tumor doubling in the treated mice relative to untreated mice as has been previously described (14) and as such accounts for differences in baseline tumor growth. Data are expressed as a mean ± SEM tumor volume. Group sizes are specified in the respective figure legends. All experimental procedures were approved in accordance with Institutional Animal Care & Use Committee (IACUC) and Yale University institutional guidelines for animal care and ethics and guidelines for the welfare and use of animals in cancer research.

### Statistical Analysis

All data are plotted as mean ± SEM unless specified. Statistical analyses were performed using GraphPad Prism 9.0 or 10.0 software (GraphPad Software Inc.), as described in the Results. P values ≤ 0.05 were considered statistically significant and all tests are two-tailed unless otherwise indicated.

## Results

### PLK4 inhibition enhances the radiation sensitivity of NSCLC *in vitro*

We defined the effects of PLK4 inhibition with CFI-400945 on NSCLC proliferation using the MTT assay. In H460 cells there was a dose-dependent reduction in proliferation with a low nanomolar (nM) IC50 (fifty percent inhibitory concentration) of 24nM (Supplemental Figure 1A). A similar effect was seen in A549 cells, where CFI-400945 inhibited proliferation in a dose-dependent manner with an IC50 of 23nM (Supplemental Figure 1B). These results are consistent with previous studies demonstrating dose-dependent inhibition of proliferation by CFI-400945 (37,44). Based upon these data we have selected a concentration of 10nM, which reduces viability by 30% and 29% in H460 and A549 respectively, for further experiments. This dose minimizes excessive toxicity to allow for the determination of radiation sensitization and decreases the potential for off-target effects known to happen at higher drug concentrations.

The effects of CFI-400945 on NSCLC radiation sensitivity were determined using the clonogenic survival assay. Consistent with short-term growth delay assays CFI400945 treatment alone reduced surviving fraction to 64% in H460 cells and 83% in A549 cells (Figures 1 A and B respectively). We found that CFI-400945 treatment resulted in significant radiosensitization of both H460 and A549 cells (Figures 1 C and D, respectively), with dose enhancement factors (DEF) at a surviving fraction of 0.2 for H460 and A549 cells were 1.6 and 1.3 respectively. We further defined the effects of the distinct PLK4 inhibitor (Centrinone) on NSCLC *in vitro* radiation sensitivity using the clonogenic survival assay. For these assays, we used sub-IC50 concentrations as determined by MTT (Supplemental Figure 1 C-D). Treatment with centrinone alone reduced the surviving fraction to 66% and 63% in H460 and A549 cells respectively. Centrinone treatment resulted in significant radiosensitization of H460 and A549 cells (Figure 1 E and F, respectively) with DEF of 2.5 and 1.4 respectively.

**Figure 1:**
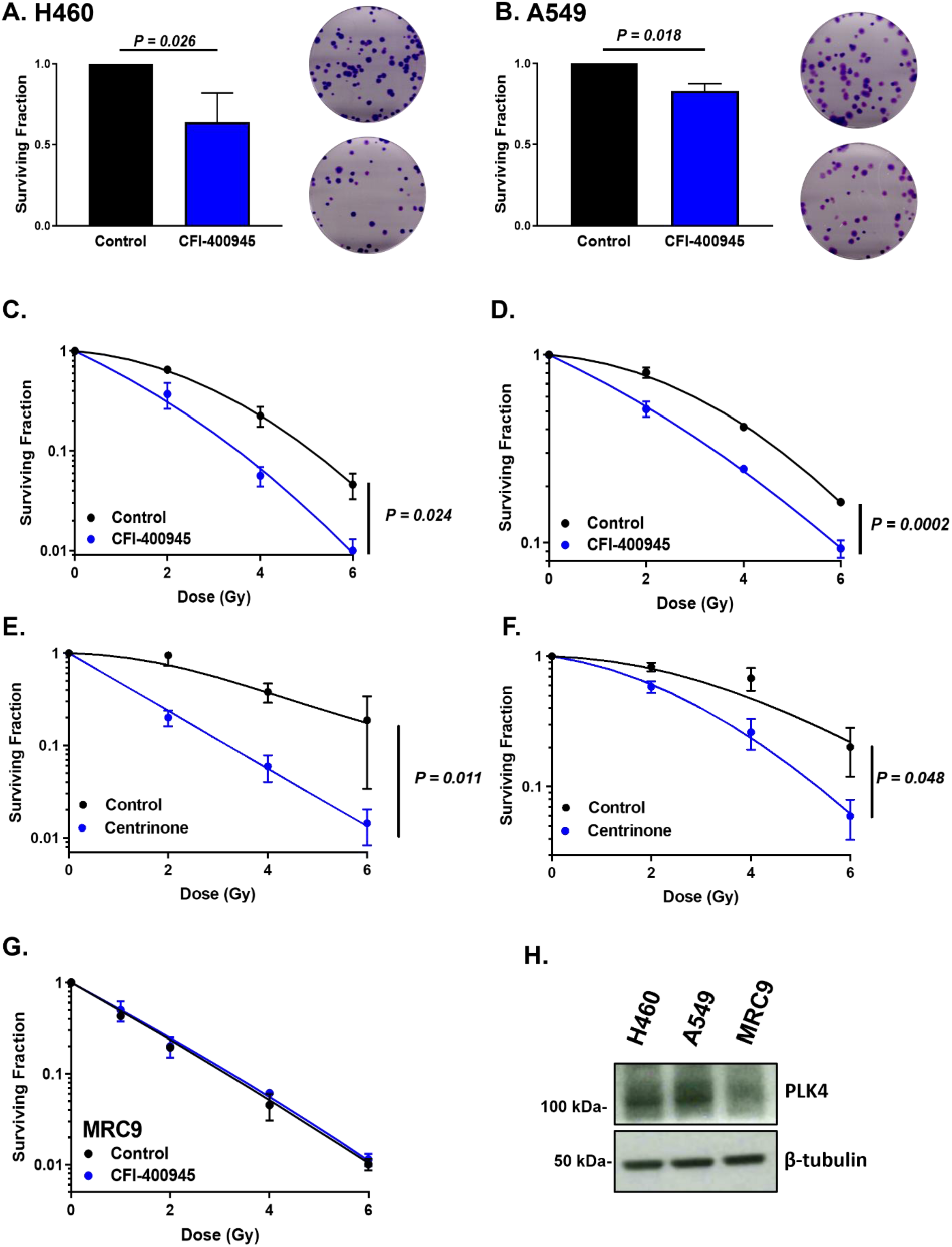
PLK4 inhibition results in tumor specific radiosensitization of NSCLC. Quantification and representative images of clonogenic survival analysis of H460 (**A**) and A549 (**B**) cells treated with CFI-400945 (10nM). Error bars represent standard error of the mean (SEM) for n=3 independent experiments analyzed by unpaired, two-tailed t-tests. Clonogenic survival analysis of H460 (**C**) or A549 (**D**) cells treated with vehicle (DMSO) or CFI-400945 (10nM) and the indicated dose of radiation. The results represent data from three independent experiments for each cell line. Data are represented as the mean ± SEM analyzed by two-way ANOVA testing. Clonogenic survival analysis of H460 (**E**) or A549 (**F**) cells treated with vehicle (DMSO) or centrinone (2.5 µM) and the indicated dose of radiation. The results represent data from three independent experiments for each cell line. Data are represented as the mean ± SEM analyzed by two-way ANOVA testing. **G.** Clonogenic survival analysis of MRC9 cells treated with vehicle (DMSO) or CFI-400945 (10nM) and the indicated dose of radiation. The results represent data from three independent experiments for each cell line. Data are represented as the mean ± SEM. **H.** Immunoblot of PLK4 expression in the indicated cell lines. Immunoblots are representative of three independent experiments.

To investigate the potential for tumor selectivity we determined the effects of CFI-400945 on the radiosensitivity of the normal lung fibroblast line, MRC9. In contrast to the 2 tumor cell lines, CFI-40095 treatment had no effects on the radiosensitivity of MRC9 cells (Figure 1G). Consistent with this tumor selectivity, MRC9 cells had lower expression of PLK4 when compared to both tumor cell lines, in line with prior analysis of lung cancer patient data (Figure 1H)(26). Overall, these data suggest that PLK4 inhibition results in tumor selective radiosensitization of NSCLC cell lines.

### CFI-400945 enhances cell-cycle arrest following radiation in lung cancer

Given our findings showing PLK4 inhibition results in tumor selective radiosensitization of NSCLC cells, we then proposed to define the mechanism of radiosensitization. Cell cycle distribution at the time of irradiation is known to be an important determinant of cellular radiosensitivity (45). Given that PLK4 is a critical regulator of centriole duplication and inhibition of PLK4 has been shown to alter cell cycle distribution we defined the effects of PLK4 inhibition using CFI-400945 on baseline cell cycle distribution after 24h of exposure (time of irradiation in clonogenic survival experiments above) (25,34,35). As shown in Figure 2A, the cell cycle distribution of H460 cells was not significantly altered at 24h after exposure to CFI-400945 as compared to vehicle treated cells. Consistently, the cell cycle distribution of A549 cells was also not significantly altered after exposure to CFI-400945 for 24h (Figure 2B). These results indicate that cell cycle redistribution into a radiosensitive phase of the cell cycle is not responsible for the observed radiosensitization with CFI-400945 treatment.

**Figure 2:**
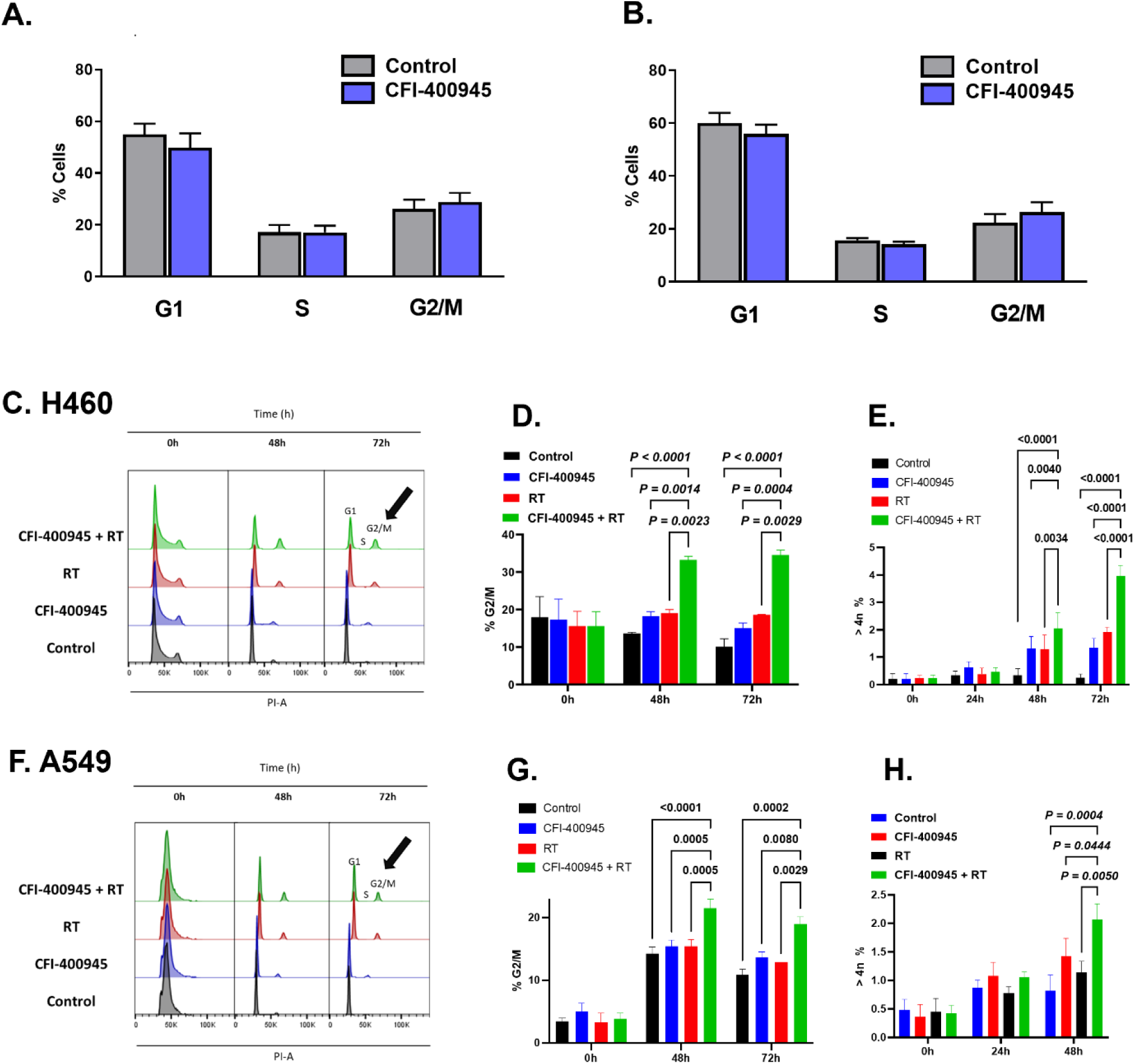
PLK4 inhibition increases radiation-induced G2/M cell cycle arrest. Cell cycle distribution of H460 (**A**) and A549 (**B**) cells treated with CFI-400945 for 24h. Error bars represent standard error of the mean (SEM) for n=3 independent experiments. Representative histogram (**C**) and quantification of G/2M (**D**) and >4N (**E**) populations in H460 cells synchronized with double thymidine block, released and treated with CFI-400945, RT (4Gy), or the combination of CFI-400945 and RT analyzed at the indicated timepoints. Representative histogram (**F**) and quantification of G/2M (**G**) and >4N (**H**) populations in A549 cells synchronized with double thymidine block, released and treated with CFI-400945, RT (4Gy), or the combination of CFI-400945 and RT analyzed at the indicated timepoints. Data are represented as the mean ± SEM from three to four independent experiments analyzed by two-way ANOVA with Fisher’s LSD post-hoc analysis without multiple comparison correction.

Cell cycle control following radiation is a critical regulator of cell survival and given the importance of PLK4 in regulating cell cycle progression we defined the effects of CFI-400945 on radiation-induced cell cycle distribution (45). Towards this end, H460 cells were synchronized using a double thymidine block, followed by the indicated treatment and analysis of cell cycle distribution (Figures 2C-D). The combination of CFI-400945 resulted in a nearly 2-fold increase in the proportion of cells in G2/M at both 48 and 72h after irradiation (Figure 2D; P<0.003). Furthermore, the combination of CFI-400945 and RT resulted in a ∼2-fold increase in the number of cells with >4n DNA content consistent with the increase in polyploid cells (P < 0.004, Figure 2E). Similar results were obtained in A549 cells where CFI-400945 treatment resulted in a 1.5-fold increase in the proportion of cells in G2/M (P< 0.001, Figure 2 F-G) and cells with >4n DNA content (P< 0.05, Figure 2H). These results suggest that CFI-400945 increases radiation-induced G2/M cell cycle arrest and increases the polyploid population consistent with decreased survival after radiation.

### CFI-400945 treatment enhances radiation-induced centrosome amplification

PLK4 is a critical regulator of centrosome amplification through the regulation of centriole biogenesis (22–24). Furthermore, DNA-damaging therapies are known to cause alterations in structure and duplication and potentiation of centrosome overduplication has been associated with increased cell death (16–19). Thus, we hypothesized that PLK4 inhibition would potentiate radiation-induced centrosome amplification. To test this hypothesis, we defined the effects of PLK4 inhibition using CFI-400945 on radiation-induced centrosome amplification using immunofluorescence microscopy in H460 cells (Figure 3A–B). Radiation and CFI-400945 monotherapy treatment resulted in moderate, time-dependent increases in the number of cells with centrosome amplification (>2 centrosome). Strikingly, the addition of CFI-400945 to radiation resulted in a nearly 2-fold increase in the number of cells with centrosome amplification when compared with radiation alone (62% vs 32%; P = 0.007). Similar findings were observed in A549 cells, where radiation and CFI-400945 monotherapy resulted in a modest increase in the number of cells with centrosome amplification, but the combination of CFI-400945 and radiation resulted in a ∼3-4-fold increase in the number of cells with centrosome amplification when compared with radiation or CFI-400945 treatment alone (Figure 3C-D; P< 0.04). These results indicate that PLK4 inhibition using CFI-400945 enhances radiation-induced centrosome amplification.

**Figure 3.**
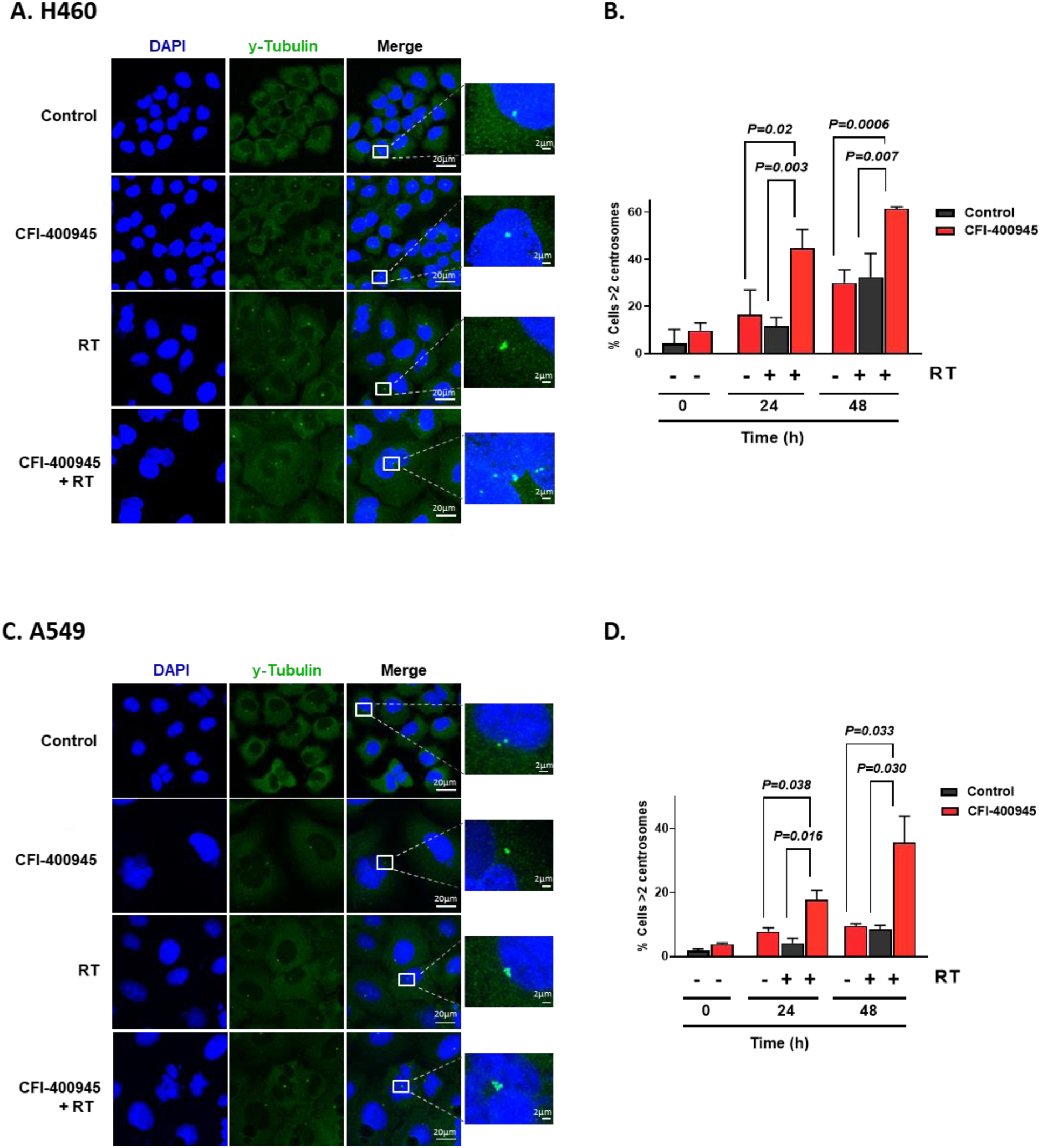
PLK4 inhibition increases radiation-induced centrosome amplification. Representative images (**A**) and quantification (**B**) of centrosome number in H460 cells treated with CFI-400945 (10nM), RT (4 Gy), or the combination of CFI-400945 and RT at the indicated time points post-treatment. Representative images (**C**) and quantification (**D**) of centrosome number in A549 cells treated with CFI-400945 (10nM), RT (4 Gy), or the combination of CFI-400945 and RT at the indicated time points post-treatment. Data are represented as the mean ± SEM from three independent experiments analyzed by unpaired, two-tailed t-tests, the scale bar is 20 µm in main images and 2 µm in insets.

### PLK4 inhibition enhances radiation-induced mitotic catastrophe

Mitotic catastrophe is the primary form of radiation-induced cell death in non-hematopoietic cells, which is evidenced by the presence of multiple distinct nuclear lobes within a cell (17,39–41,43,46). Additionally, radiation-induced mitotic catastrophe has been associated with overduplication of centrosomes (16–19). Thus, given our data showing a significant increase in radiation-induced centrosome amplification by CFI-400945, we hypothesized that CFI-400945 would result in an increase in cell death by mitotic catastrophe and therefore examined the effects of CFI-400945 on markers of mitotic catastrophe after radiation exposure. Indeed, CFI-400945 treatment resulted in a 3-4-fold increase in the proportion of H460 cells undergoing radiation-induced mitotic catastrophe number (12.0% vs 47.5% at 48h; P< 0.0001, Figure 4A-B). Similar findings were seen in A549 cells, where PLK4 inhibition using CFI-400945 resulted in a ∼2-fold increase in the proportion of cells undergoing radiation-induced mitotic catastrophe (20.1% vs 45.7% at 48h, P=0.002, Figure 4C-D). Together these results suggest that PLK4 inhibition enhances radiation-induced centrosome amplification, which then contributes to an increase in the number of cells undergoing mitotic catastrophe.

**Figure 4:**
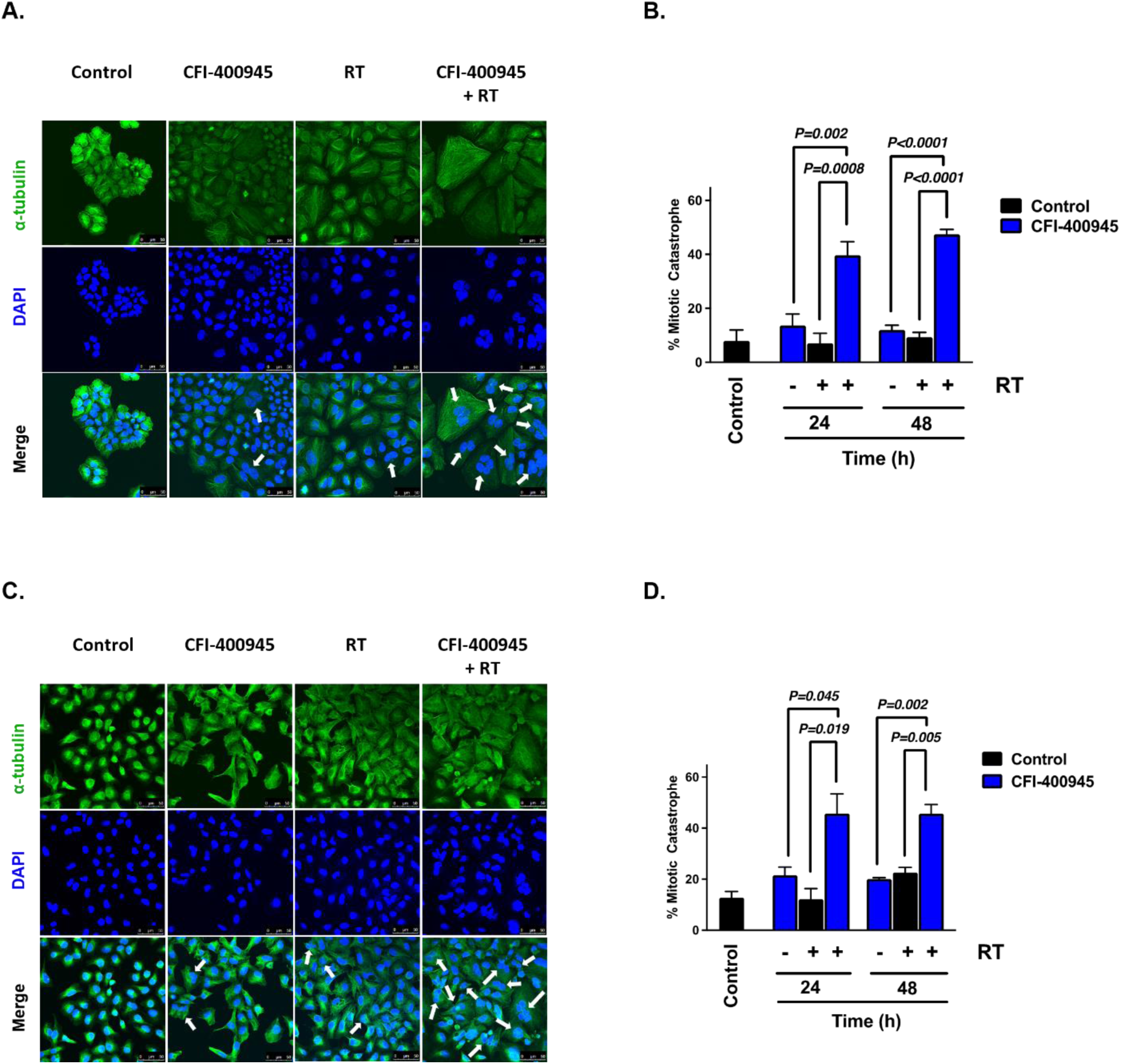
PLK4 inhibition enhances radiation-induced mitotic catastrophe. Representative images (**A**) and quantification (**B**) of mitotic catastrophe in H460 cells treated with CFI-400945 (10nM), RT (2 Gy), or the combination of CFI-400945 and RT at the indicated timepoints post-treatment. Representative images (**C**) and quantification (**D**) of mitotic catastrophe in A549 cells treated with CFI-400945 (10nM), RT (2 Gy), or the combination of CFI-400945 and RT at the indicated timepoints post-treatment. Arrows highlight examples of cells with multiple distinct nuclear lobes, a marker of mitotic catastrophe. Data are represented as the mean ± SEM from three independent experiments analyzed by unpaired, two-tailed t-tests.

### CFI-400945 enhances the radiation response of NSCLC *in vivo*

To determine whether the enhancement of radiosensitivity with CFI-400945 observed *in vitro* translated to *in vivo* NSCLC tumor xenograft models, we defined the effects of CFI-400945 treatment on tumor growth delay in NSCLC tumor xenograft models. Towards this end, mice bearing H460 tumor xenografts were randomized to receive vehicle, CFI-400945 (7.5 mg/kg daily, oral gavage), radiation (2 Gy x 5 daily fractions), or the combination of radiation and CFI-400945 as depicted in Figure 5A. Radiation treatment or CFI-400945 had modest effects on tumor growth, however, the combination of radiation and CFI-400945 significantly inhibited tumor growth when compared to monotherapy treatment alone (Figure 5B). Tumor growth delay, calculated as the time to reach a volume of 400mm^3^ (i.e. tumor doubling) was ∼3-6-fold longer with the combination of radiation and CFI-400945 than with radiation or CFI-400945 monotherapy alone (5.4 days vs 1.7 days vs 0.1 days respectively, P< 0.0001; Figure 5C). Furthermore, the median time to tumor doubling was significantly increased for combination treatment (8.3 days) compared with radiation (5.1 days) or CFI-400945 (2.9 days) (Figure 5D). From a toxicity perspective treatment was well tolerated with a transient <5% weight loss associated with CFI-400945 administration (Suppl Figure 3A-B). We also tested the effects of CFI-400945 on tumor growth in mice bearing A549 tumor xenografts. Consistent with the results from H460 tumor xenografts, the combination of radiation and CFI-400945 treatment resulted in significant inhibition of tumor growth with a ∼2-3-fold increase in tumor growth delay when compared to RT or CFI-400945 treatment alone (13.2 days vs 7.1 days vs 3.4 days, respectively, P< 0.03; Figure 5E-F). Lastly, the median time to tumor doubling was significantly longer with combination treatment (22.4 days) as compared with CFI-400945 (10.1 days) or radiation (17.0 days) (Figure 5G). These results indicate that the combination of CFI-400945 and radiation enhances NSCLC tumor radiation sensitivity.

**Figure 5:**
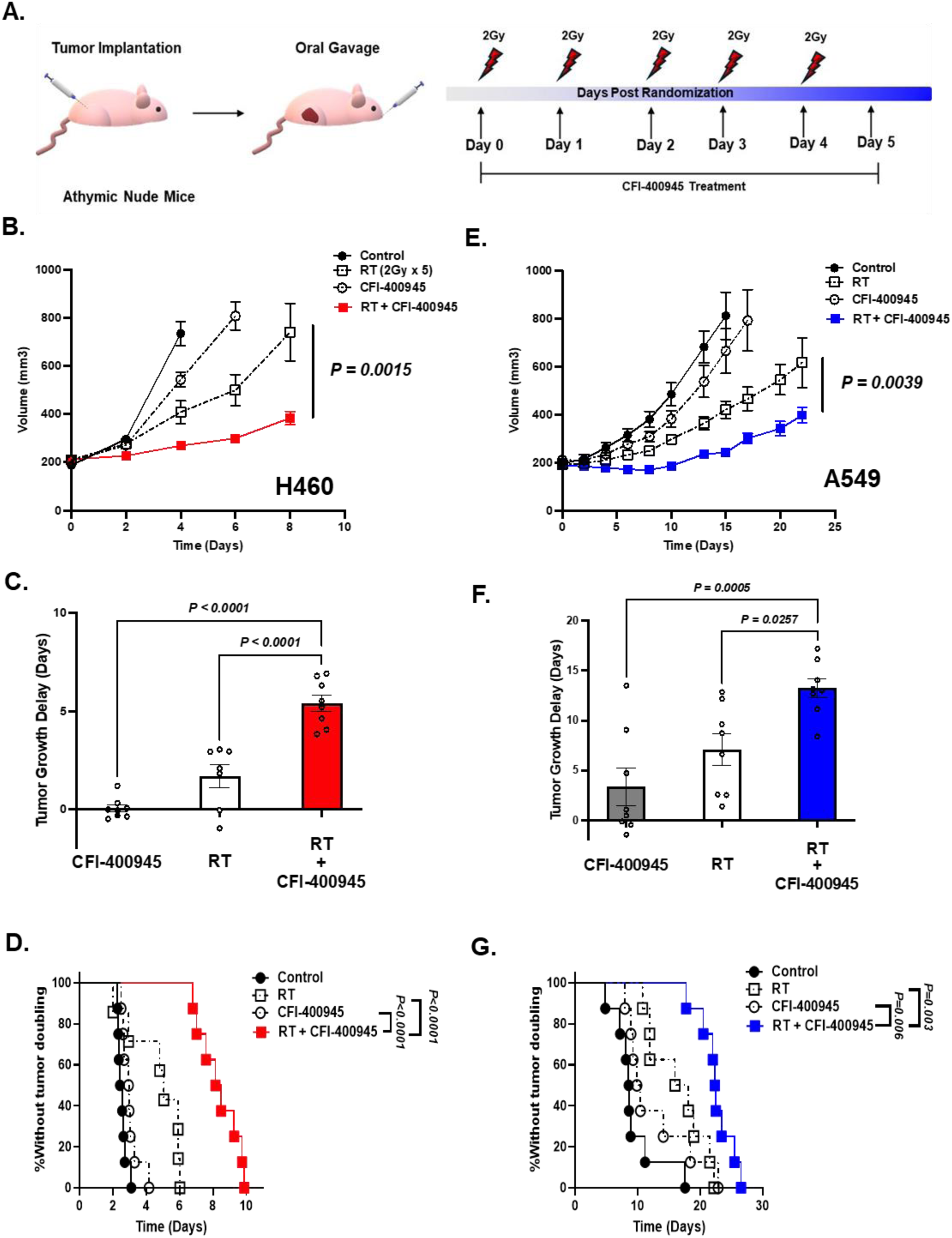
PLK4 inhibition with CFI-400945 enhances *in vivo* NSCLC response to radiation therapy. **A.** Schematic of *in vivo* tumor growth delay experiments and fractionated treatment protocol with radiation and CFI-400945 treatment. **B.** Tumor growth curves for H460 cells implanted subcutaneously in athymic nude mice with the indicated treatment. *P*-value from two-way ANOVA testing. **C.** Quantification of tumor growth delay from the indicated treatment (time to reach 400mm^3^) from **B.** Statistical analysis was performed with one-way ANOVA testing with Tukey’s post-hoc multiple comparison correction. **D.** Kaplan–Meier curves of percent tumors without tumor doubling from **B**. Statistical analysis was performed by log-rank testing. Error bars in **B-D** represent the SEM of n=8 mice for CFI-400945, CFI-400945 + RT, n = 9 mice for vehicle group, and n = 7 mice for RT. **E.** Tumor growth curves for A549 cells implanted subcutaneously in athymic nude mice with the indicated treatment. P-value from two-way ANOVA testing. **F.** Quantification of tumor growth delay from the indicated treatment (time to reach 400mm^3^) from **E.** Statistical analysis was performed with one-way ANOVA testing with Tukey’s post-hoc multiple comparison correction. **G.** Kaplan–Meier curves of percent tumors without tumor doubling from **E.** Statistical analysis was performed by log-rank testing. Error bars in **E-G** represent the SEM of n=8 mice.

## Discussion

PLK4 is a critical regulator of centrosome amplification through the regulation of centriole duplication (22–24). Aberrant PLK4 expression is a known feature of NSCLC and other malignancies and inhibition of PLK4 activity results in tumor cell death (25,26,33,35). As such, PLK4 has emerged as a therapeutic target for multiple cancer types with the development of multiple PLK4 inhibitors (24,25,29). CFI-400945 is a first-in-class, orally available PLK4 inhibitor that is undergoing clinical trial evaluation and has FDA orphan drug designation in the treatment of acute myelogenous leukemia (25,26,29,33,35,47,48). In the context of DNA damage, there is close coordination of centrosome duplication and the DNA damage response that is necessary to prevent abnormal centrosome amplification, genomic instability, and cell death. In the work presented here, we show that PLK4 inhibition, using the clinically available CFI-400945 compound, enhances the radiation sensitivity of NSCLC cell lines *in vitro*, but has no effects on the radiation sensitivity of the normal lung fibroblast cell line, MRC9. We show that CFI-400945 precipitates sustained G2/M arrest after radiation, enhances radiation-induced centrosome overamplification, and ultimately results in increased cell death through mitotic catastrophe. Finally, we show that PLK4 inhibition using CFI-400945 enhances the radiation sensitivity of NSCLC tumor xenografts. Together these results identify PLK4 inhibition, using the clinically available CFI-400945 molecule, as a potential strategy to enhance the radiation response of NSCLC, a treatment refractory disease with a high propensity for local recurrence after conventional radiation therapy.

Close coordination of centrosome duplication and the DNA damage response is necessary to prevent abnormal centrosome amplification, mitotic errors, and subsequent cell death (12,49,50). In this context ionizing radiation has been shown to increase centrosome amplification and subsequent mitotic failure (16,51–53). PLK4 has a critical role in the regulation of centrosome amplification through control of centriole biogenesis, mitotic fidelity, and thus maintenance of genome stability (23–25). Our data show that the mechanism of radiosensitization, using low nanomolar doses of CFI-400945, involves potentiation of radiation-induced centrosome amplification and increased G2/M cell cycle arrest, rather than alterations of cell cycle distribution prior to irradiation. We further show that this treatment led to increased cell death through mitotic catastrophe, the main mechanism of cell death induced by radiation in non-hematopoietic tumors. Our findings are supported by data demonstrating that cells with supernumerary levels of centrosomes (>2 centrosomes) are prone to errors in mitosis leading to increased cell death by mitotic catastrophe and are consistent with a recent report showing increased radiation-induced centriole amplification using CFI-400945 in breast cancer cell lines (37). Collectively our data support a role for PLK4 in regulating radiation-induced centrosome amplification whereby inhibition of PLK4 using CFI-400945 potentiates radiation-induced centrosome amplification in NSCLC and subsequent cell death through mitotic catastrophe.

We show that PLK4 inhibition by CFI-400945 or Centrinone results in the radiosensitization of NSCLC *in vitro* and *in vivo*. A critical characteristic of a novel radiation sensitizing agent is the ability to selectively enhance the radiation sensitivity of tumor cells over normal tissue, thus maintaining a relevant therapeutic window. This is of crucial importance in NSCLC given the deleterious effects of non-selective dose escalation leading to worsened outcomes likely secondary to normal tissue toxicity as illustrated in the RTOG0617 Phase III randomized trial (5,54,55). In this work we show that PLK4 inhibition has no effects on the radiation sensitivity of the normal lung fibroblast cell line, MRC9, providing initial preclinical evidence that PLK4 may be a tumor selective target for radiosensitization. These data are consistent with the known overexpression of PLK4 in clinical NSCLC patient specimens when compared with surrounding normal tissue, as well as our data showing higher expression of PLK4 in NSCLC tumor cell lines when compared with normal lung fibroblasts (26,56). Further supporting the notion of tumor selectivity by the observation that PLK4 overexpression-induced centrosome amplification leads to aneuploidy, tumorigenesis, and worse oncologic outcomes, thus potentially identifying a form of oncogene addiction in tumors (25–27). Overall, our data support the role of PLK4 as a tumor selective target for radiosensitization in NSCLC.

We have used 10nM CFI-400945 for our experiments, a concentration that has been shown to have partial inhibitory function of PLK4 and thereby suppresses PLK4 autophosphorylation, leading to centriole and centrosome overduplication, as opposed to full PLK4 inhibition leading to centrosome depletion (33,35). Our findings presented here are consistent with this bimodal inhibition of PLK4, whereby we show increased centrosome amplification using low nanomolar concentrations of CFI-400945 that then leads to increased cell death through mitotic catastrophe. While off-target effects of CFI-00945 on aurora B have been reported, the 10nM concentration of CFI-400945 we used is significantly below the IC50 for off-target aurora B inhibition (98nM) and this concentration has previously been shown to have no effects on aurora B signaling thus minimizing off-target effects (33,35). This is supported by our experiments showing similar degrees of radiosensitization with Centrinone, a chemically distinct PLK4 inhibitor. Thus, while off-target effects of CFI-400945 have not been fully ruled out, our data support further evaluation of the clinically available PLK4 inhibitor CFI-400945 in combination with radiation in NSCLC.

In summary, we show that the use of the first-in-class, clinically available PLK4 inhibitor CFI-400945 enhances the radiation sensitivity of NSCLC both *in vitro* and *in vivo* through increased centrosome amplification and enhanced cell death through mitotic catastrophe. In contrast, CFI-400945 has no effects on the radiosensitivity of normal lung fibroblasts, suggesting tumor specific radiosensitization, thus improving the therapeutic ratio. Given the known resistance of NSCLC to standard-of-care, curative-intent radiation therapy, these results suggest that further translational and clinical evaluation of CFI-400945 and radiation may be a relevant approach to improve outcomes in NSCLC.

## Acknowledgments

We thank the Yale West Campus Imaging Core for the support and assistance in this work. This research utilized the services of the Irradiator Shared Resource through the Yale Cancer Center Support Grant (P30-CA016359). We thank Yale Flow Cytometry for their assistance with and use of the LSR II flow cytometer. The Flow Cytometry Core is supported in part by an NCI Cancer Center Support Grant # NIH P30CA016359.

## Data Availability Statement

Data associated with this study are available by reasonable request to the corresponding author.

## Author contributions

I. Dominguez-Vigil: Conceptualization, data curation, formal analysis, investigation, methodology, writing–original draft, writing–review, and editing.

K. Banik: Conceptualization, data curation, formal analysis, investigation, methodology, writing– original draft, writing– original draft, writing–review and editing.

M. Baro: Acquisition of data, analysis and interpretation, writing–review and editing.

J. N. Contessa: Conceptualization, resources, writing – review and editing.

T. J. Hayman: Conceptualization, data curation, resources, formal analysis, investigation, supervision, project administration, writing– original draft, review and editing.

