## Supplementary Data for "PLK4 inhibition as a strategy to enhance non-small cell lung cancer radiosensitivity"

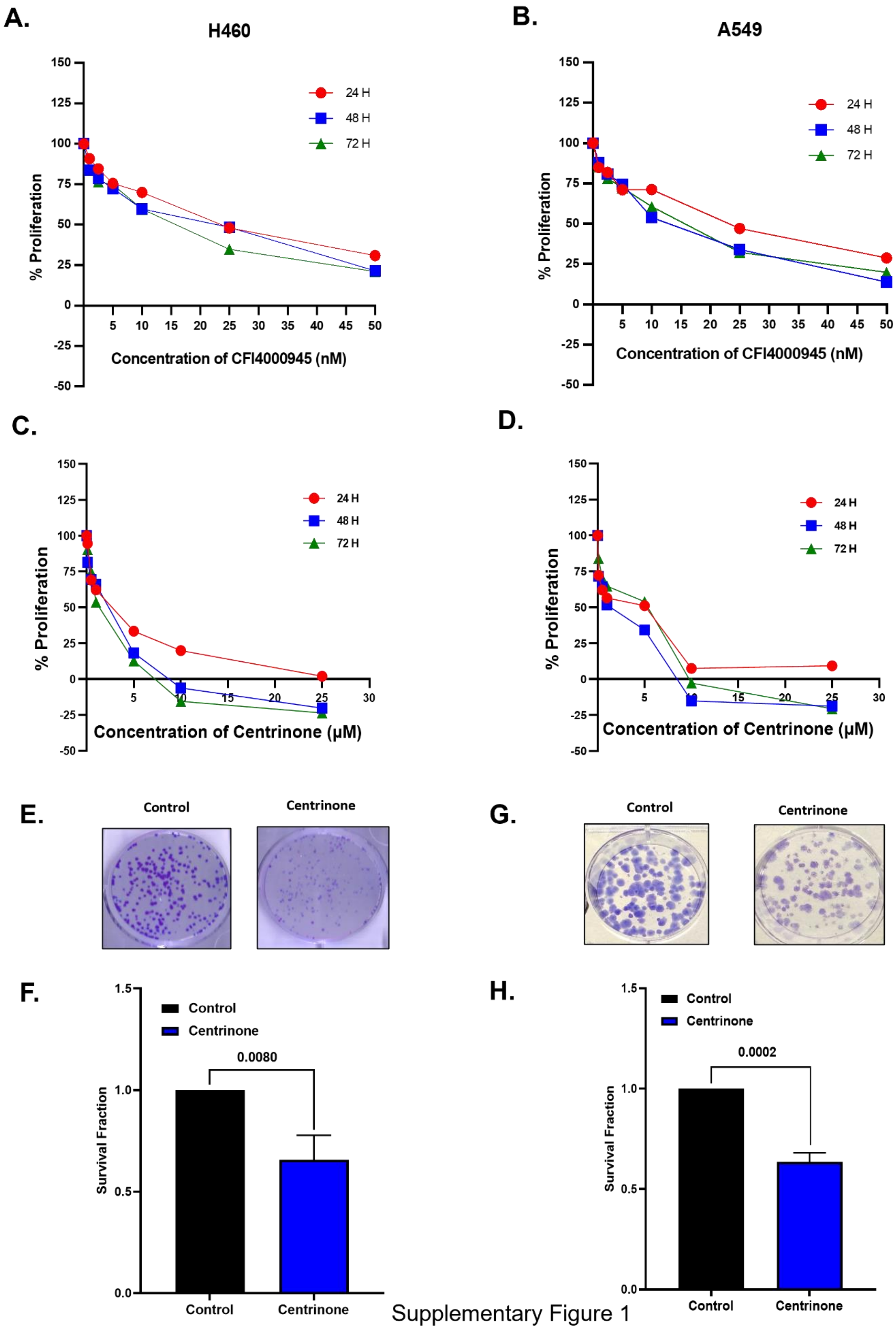


Supplementary Figure 1

**Supplemental Figure 1. Plk4 inhibition results in a dose-dependent reduction of NSCLC proliferation.** Quantification of the effects of CFI-400945 on H460 (**A**) and A549 (**B**) proliferation treated for the indicated duration of time. Error bars represent SEM for n=3 independent experiments. Quantification of the effects of Centrinone on H460 (**C**) and A549 (**D**) proliferation treated for the indicated duration of time. Error bars represent SEM for n=3 independent experiments. Representative images (**E**) and quantification (**F**) of clonogenic survival analyses of H460 cells treated with Centrinone (2.5 μM). Error bars represent SEM for n=3 independent experiments analyzed by unpaired, two-tailed t-tests. Representative images (**G**) and quantification (**H**) of clonogenic survival analyses of A549 cells treated with Centrinone (2.5 μM) Error bars represent SEM for n=3 independent experiments analyzed by unpaired, two-tailed t-tests.

**A)**

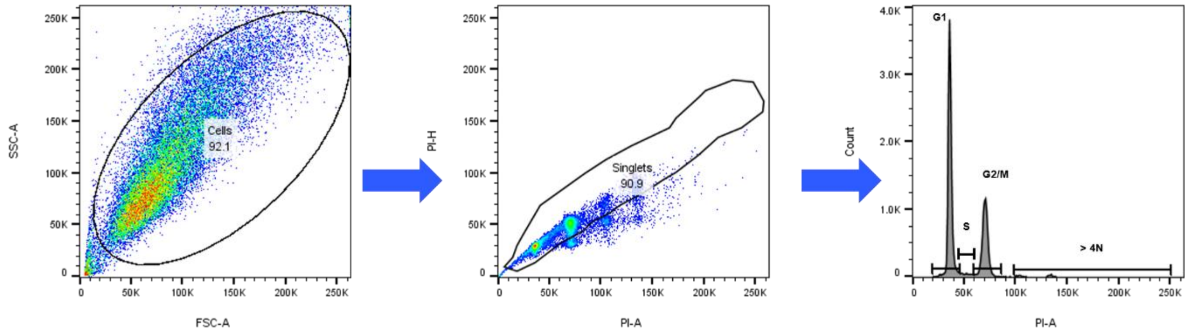


Supplementary Figure 2

**Supplementary Figure 2**. **Flow cytometry gating strategy.** (**A)** Gating strategy for DNA content cell cycle analysis. Events were plotted by forward and side scatter parameters (FSCA-A vs SSC-A), then doublets were discriminated (PI-H vs PI-A), and G1, S, G2/M, and >4N populations were determined.

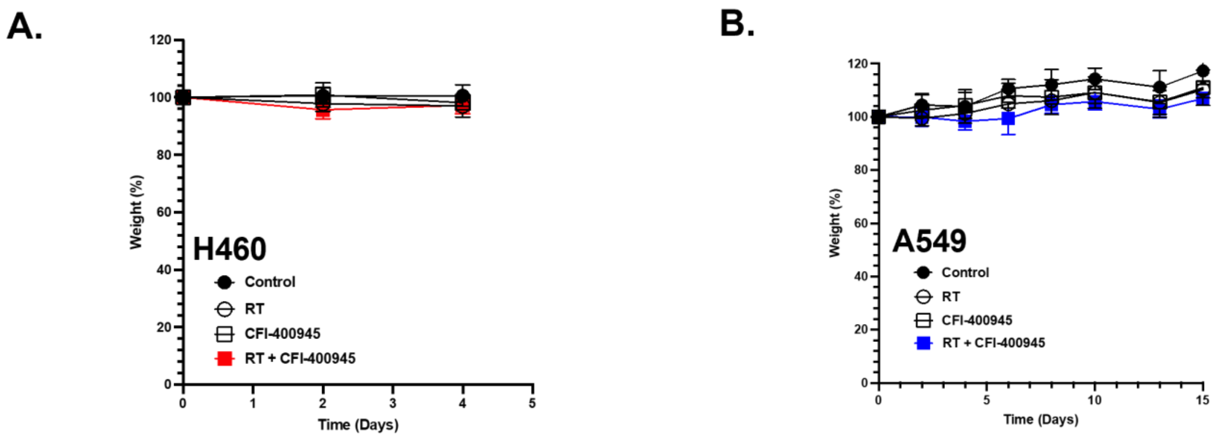


Supplementary Figure 3

**Supplemental Figure 3. CFI-400945 treatment does not lead to significant weight loss in combination with radiation. A.** Weight as stratified by treatment group for H460 tumor growth delay experiment from Figure 5B. **B.** Weight as stratified by treatment group for A549 tumor growth delay experiment from Figure 5E.
